# Evolution of compound eye morphology underlies differences in vision between closely related *Drosophila* species

**DOI:** 10.1101/2023.07.16.549164

**Authors:** Alexandra D Buffry, John P Currea, Franziska A Franke, Ravindra Palavalli-Nettimi, Andrew J Bodey, Christoph Rau, Nazanin Samadi, Stefan J Gstöhl, Christian M Schlepütz, Alistair P McGregor, Lauren Sumner-Rooney, Jamie Theobald, Maike Kittelmann

## Abstract

Insects have evolved complex visual systems and display an astonishing range of adaptations for diverse ecological niches. Differences in eye size within and between *Drosophila* species provide the opportunity to study the impact of eye structure on vision. Here we further explored differences in *D. mauritiana* and its sibling species *D. simulans* and confirmed that *D. mauritiana* have rapidly evolved larger eyes as a result of more and wider ommatidia than *D. simulans* since their recent common ancestor. The functional impact of eye size, and specifically ommatidia size, is often only estimated based on the rigid surface morphology of the compound eye. Therefore, we used 3D synchrotron radiation tomography to measure optical parameters in 3D, predict optical capacity, and compare the modelled vision to *in vivo* optomotor responses. Our optical models predicted higher contrast sensitivity for *D. mauritiana*, which we verified by presenting sinusoidal gratings to tethered flies in a flight arena. Similarly, we confirmed the higher spatial acuity predicted for *Drosophila simulans* with smaller ommatidia and found evidence for higher temporal resolution.

## Introduction

Insect compound eyes exhibit remarkable variation in size as a result of differences in the number and diameter of the individual subunits, known as ommatidia. For example, the silverfish *Tricholepidion gertschi has* ∼40 ommatidia (Blanke et al., 2014) whereas dragon- and damselflies (Odonata), sport up to 30,000 ommatidia (Sherk, 1978). Differences in eye size as well as the number, size, and angles between facets allow different visual behaviours, lifestyles and adaptation to a large range of environments (e.g. Land, 1999, 2012; Land et al., 1999; Greiner, Ribi and Warrant, 2004; Meyer-Rochow and Mishra, 2007; Gonzalez-Bellido, Wardill and Juusola, 2011; Nilsson, 2013; Tocco, Dacke and Byrne, 2019; Meece, Rathore and Buschbeck, 2021; Johnson and Rutowski, 2022; Pichaud and Casares, 2022). How these diverse eyes of insects have evolved and to what extent even small changes in the optics affect vision is still not well understood. Investigating and comparing natural variation in eye size and composition, and its impact on optical capacity within and between closely related species can provide valuable insight into the functional evolution of the insect eye. Generally, wider ommatidia can harvest more light, allowing greater sensitivity, while more ommatidia and narrower inter-ommatidial angles can provide greater acuity (Snyder, Stavenga and Laughlin, 1977; Land, 1997; Land and Nilsson, 2012). Ommatidia diameter and number therefore represent a trade-off that is optimised for the specific visual needs of each insect species, strain, sex or morph.

Several studies have reported the extensive variation in ommatidia number, ommatidia diameter, and overall eye size within and between species of *Drosophila* (Posnien et al., 2012; Arif et al., 2013; Hilbrant et al., 2014; Keesey et al., 2019; Ramaekers et al., 2019; Gaspar et al., 2020). There is evidence that some of this is the result of trade-off between eye size and antennal size and, by extension, the visual and olfactory systems, as well as overall head capsule size (Keesey et al., 2019; Ramaekers et al., 2019; Özer and Carle, 2020). The genetic basis of these differences in eye size is complex but, in some cases, the underlying genes and developmental mechanisms have been characterised (Arif et al., 2013; Norry and Gomez, 2017; Ramaekers et al., 2019; Reis et al., 2020; Buchberger et al., 2021; Torres-Oliva et al., 2021). We previously found that one *D. mauritiana* strain has larger eyes than its sibling species *D. simulans* as a result of wider ommatidia, potentially caused by differential expression of the transcription factor Orthodenticle (Otd) during eye development (Arif et al., 2013; Gaspar et al., 2020; Torres-Oliva et al., 2021). The larger eyes of *D. mauritiana* with respect to *D. simulans* are also associated with reciprocal changes in the distance between the eyes (face width), but the antennae were not examined (Arif et al., 2013).

Optical parameters can also vary within eyes: killer flies, including *Coenosia attenuata*, have evolved specialised wide frontal ommatidia with small interommatidial angles for diurnal aerial hunting (Gonzalez-Bellido, Wardill and Juusola, 2011), and males of many families of dipterans have enlarged ommatidia in the dorso-frontal region of the eye that allows them to detect and pursue females in flight (Perry and Desplan, 2016). In *D. mauritiana* and *D. simulans* also exhibit structural variation across the eye, with significantly wider anterior frontal ommatidia than central and posterior ommatidia in both males and females (Torres-Oliva et al., 2021).

However, to fully understand the functional impact of eye size variation within and between *Drosophila* species it is crucial to test predictions based on eye morphology *in vivo*. Here we further explored the variation in ommatidia number and diameter that contribute to eye size differences in males and females among different *D. mauritiana* and *D. simulans* strains. We used synchrotron radiation microtomography to obtain 3D information of optical parameters in focal strains of these two species and tested predicted differences in their vision via optomotor responses in a virtual reality flight simulator *in vivo*.

## Results

### Ommatidia number and size vary within and between closely related *Drosophila* species

The overall eye size of compound eyes is determined by ommatidia number and size (here reflected by facet area). While *D. mauritiana* generally have larger eyes than the closely related species *D. simulans* (Hämmerle and Ferrús, 2003; Posnien *et al*., 2012; Gaspar *et al*., 2020), it remains unclear if this difference is always caused by one or both parameters. We analysed total eye area, central facet area and ommatidia number from scanning electron microscopy images in multiple strains of both species and found a negative correlation between central ommatidia facet size and number in *D. simulans* (Fig. 1, Suppl. Fig. 1) suggesting a potential trade-off between these characteristics. In contrast, *D. mauritiana* had generally wider and more numerous ommatidia and consequently overall larger eyes than *D. simulans* and the trade-off seen in *D. simulans* was absent in females. Interestingly, *D. mauritiana* males showed a positive correlation between ommatidia number and facet size and (Fig. 1, Suppl. Fig. 1).

**Figure 1:**
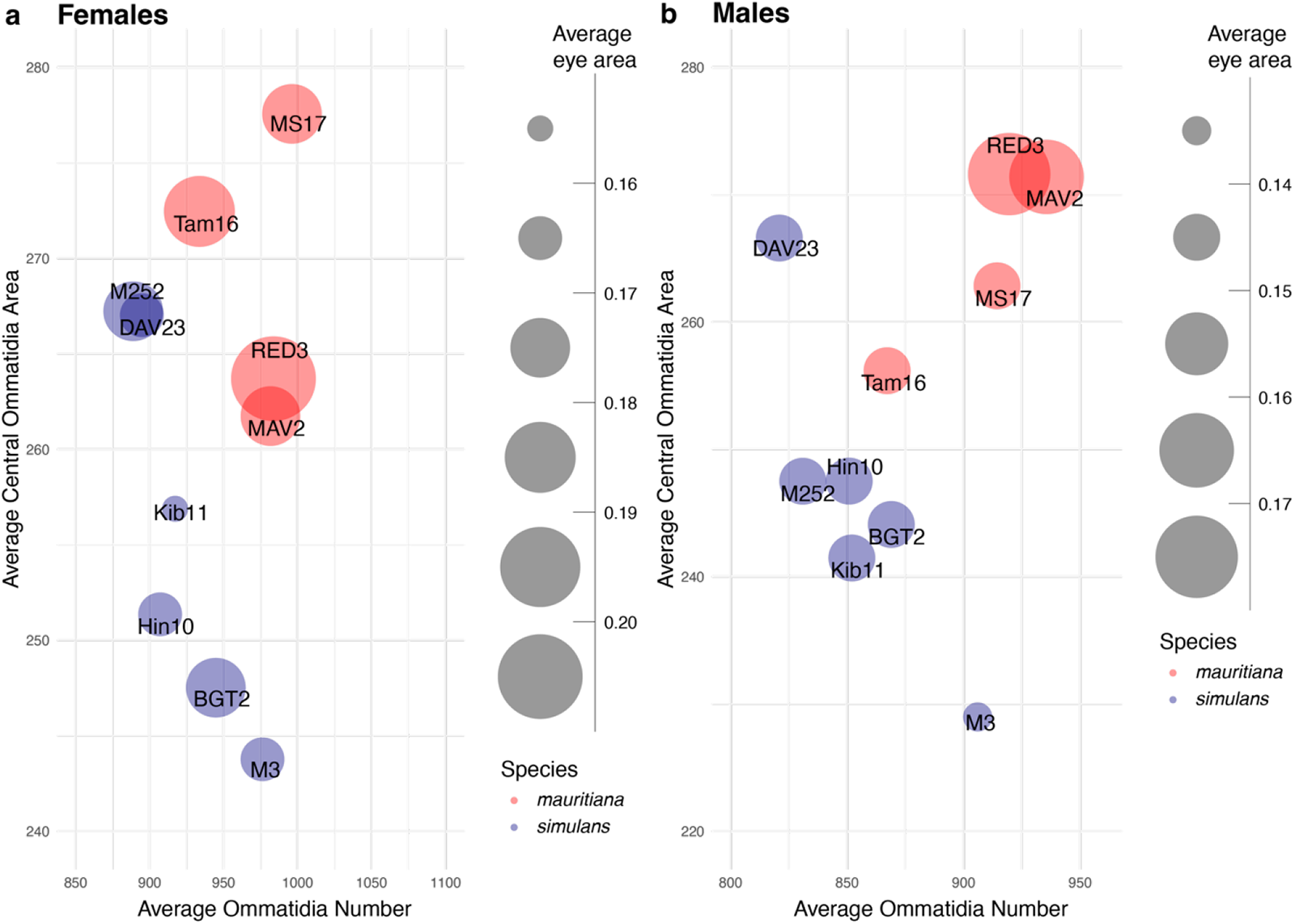
Variation in eye size, ommatidia number and ommatidia size across closely related *D. mauritiana* and *D. simulans*. Average eye size (mm^2^, circle area) of *D. simulans* (blue) and *D. mauritiana* (red) strains (circle labels) plotted against ommatidia number and ommatidia facet area (in μm^2^) for females (a) and males (b). *D. mauritiana* generally have larger eyes with more and larger ommatidia compared to *D. simulans*. Eye size was measured from side view scanning electron micrographs of single eyes.

To test whether larger eyes of *D. mauritiana* were an effect of overall larger body size we also measured second-leg tibia length, which have been previously used as a proxy for body size (Sokoloff, 1966; Posnien et al., 2012; Krause et al., 2019) and the length of the L3 wing vein as an estimate of overall wing size (Lack *et al*., 2016). The tibia of *D. mauritiana* were not generally larger than tibia of *D. simulans* (Suppl. Fig. 2), suggesting the increase in eye size has evolved independently of body size (Hämmerle and Ferrús, 2003; Posnien et al., 2012; Arif et al., 2013). Consistent with this, tibia size was only positively correlated with eye size in a subset of strains in both species (Suppl. Fig. 3). Interestingly, wing size is generally smaller in *D. mauritiana* strains, and we found strain-specific positive, negative or no correlation with eye size (Suppl. Fig. 4).

While some *D. simulans* and *D. mauritiana* strains overlap in either ommatidia area or number, none of the strains overlapped in both parameters, leading to the clear separation of the species in eye composition (Fig. 1). Previously a large-effect quantitative trait locus has been identified that explains about 30% of the eye size difference between *D. simulans* and *D. mauritiana* (Arif et al., 2013; Torres-Oliva et al., 2021) due to differences in ommatidia area. However, the functional consequences for vision in these flies remain unknown. To test this, we selected strains *D. simulans* M3 and *D. mauritiana* RED3 which have very similar ommatidia numbers but significantly different mean ommatidia (facet) areas (Fig. 1). We first performed detailed 2D and 3D morphological analysis of optical parameters of these focal strains to model their vision, and subsequently tested our predictions with behavioural experiments.

### Facet size and shape change in a dorsal to ventral-anterior gradient across species and sexes

*Drosophila* compound eyes are 3D structures that are roughly shaped like a hemisphere. To analyse optical parameters across the entire eye, we used synchrotron radiation microtomography to collect high-resolution 3D image data of entire eyes and associated brain structures in *D. simulans* M3 and *D. mauritiana* RED3 (Fig. 2). Automated segmentation and measurement of individual facets for three individuals of each species and sex revealed a size gradient from smaller dorsal-posterior to larger anterior-ventral facets in both focal strains and both sexes (Fig. 2a). Facet size was overall smaller in *D. simulans* M3 and the size difference between females and males was more pronounced in *D. simulans* M3 compared to *D. mauritiana* RED3 (Fig. 2a). Analysis of ommatidia numbers and neuropil volumes indicates that lamina and medulla are proportionally scaled down with ommatidia number in males (Fig. 2b-d).

Additionally, we used geometric morphometric analysis of facet shapes to compare central ommatidia to frontal ommatidia in both sexes of *D. simulans* M3 and *D. mauritiana* RED3 (Fig. 2e and f): the six corners of each facet were landmarked and analysed via principal component and hierarchical clustering analysis. We recovered three cluster, which can be interpreted as three distinct facet shapes. Clusters 1 and 2 contained only frontal lenses, and cluster 3 contained only central lenses indicating that the position of the facet within the eye influences lens shape. Frontal lenses (clusters 1+2) were defined by longer dorsal and ventral edges (PC1 = 87.3% variation) than central lenses (cluster 3). Within the frontal lenses, PC2 (4.5% variation) and PC9 (<1% variation) separated clusters 1 and 2, with cluster 1 being slightly elongated along the antero-posterior axis. There were no differences in sex (Chi sq=0.07, df=2, p=0.967) or strain (Chi sq=2.74, df=2, p=0.254) between clusters, implying that these factors do not influence facet shape.

**Figure 2:**
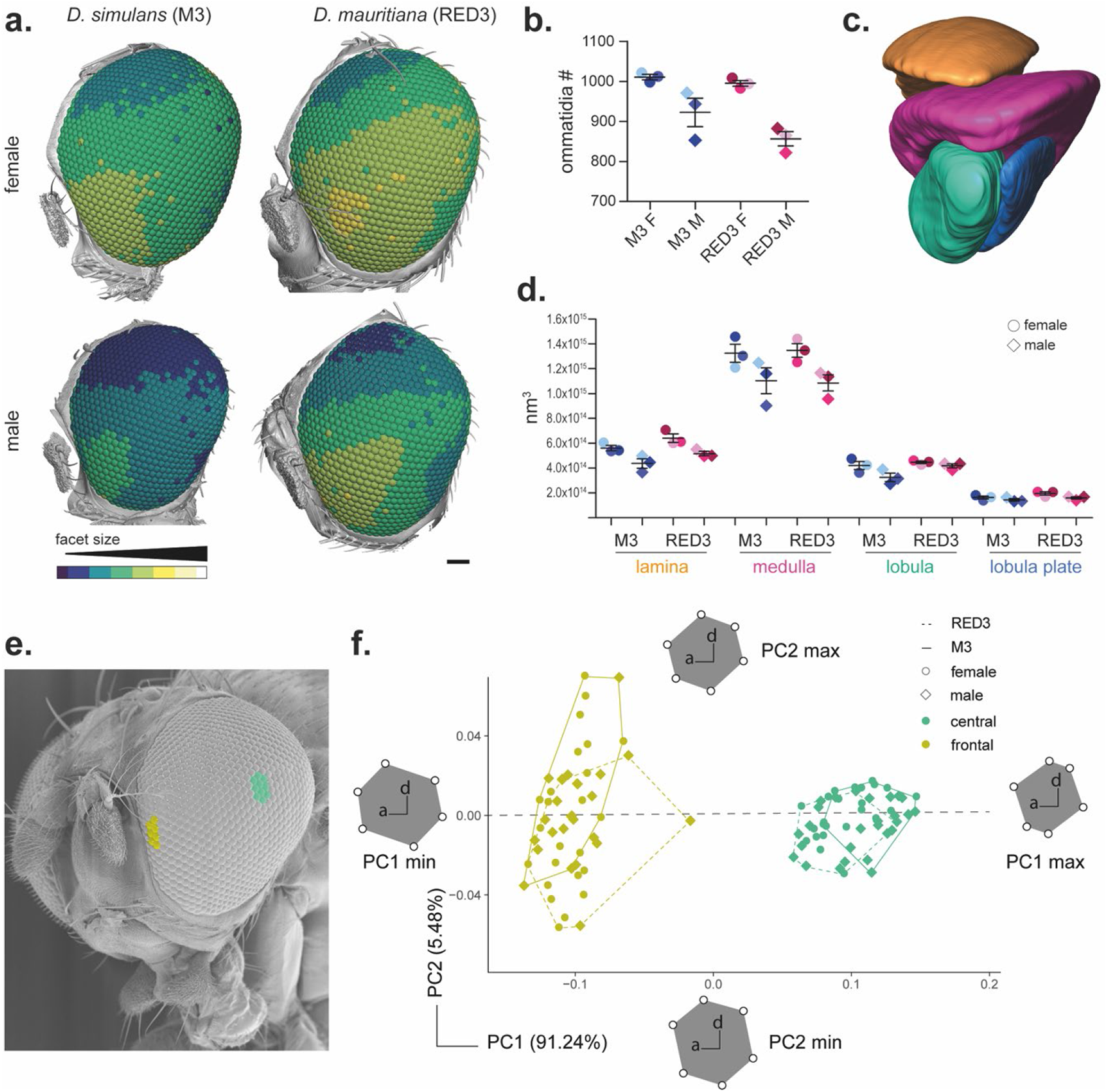
3D analysis of ommatidia size and shape in male and female *D. simulans* M3 and *D. mauritiana* RED3. **a.** synchrotron radiation microtomography analyses of males and females show a gradient from smaller to larger ommatidia from dorsal to anterior-ventral. *D. simulans* M3 females and especially males show an overall shift to smaller ommatidia compared to *D. mauritiana* RED3. **b.** Analysis of ommatidia numbers in three individuals per species and sex (including a.) show similar ommatidia numbers in females of both species and fewer ommatidia in males in line with overall smaller eye size. **c.** Segmentations of optic lobes (lamina, medulla, lobula and lobula plate) corresponding to male and female *D. simulans* and *D. mauritiana* in c. were used for volume analysis **c.** Optic lobes size scales with ommatidia number (b.): males of *D. simulans* M3 *and D. mauritiana* RED3 have generally smaller neuropils, most evident in lamina and medulla. **e – f.** Shape analysis of frontal and central ommatidia (as indicated) reveals separate clustering of frontal and central ommatidia for both species.

### *D. mauritiana* RED3 has greater optical sensitivity than *D. simulans* M3, especially in the frontal and ventral visual field

To compare the optical capacity of both fly strains and their variation across the visual field, we implemented the open-source Python-based automated pipeline ODA (Currea et al., 2023) which estimates the location and approximate orientation of each lens with high resolution across the eye. This generated eye maps of the volume (Suppl. Fig 5a), diameter (Suppl. Fig 5b), cross-sectional area (Suppl. Fig 5c), and length (Suppl. Fig 6a) of the corneal lenses of the ommatidia and the mean IO angle of each lens with its nearest neighbours (Suppl. Fig 6b). Three male and three female eyes were scanned from both *D. simulans* M3 and *D. mauritiana* RED3. The coordinates were rotated manually to align the eye equators horizontally, visible as a horizontal band of smaller ommatidia in Figure 3 a (and Suppl. Fig. 6a, b, and c).

**Figure 3:**
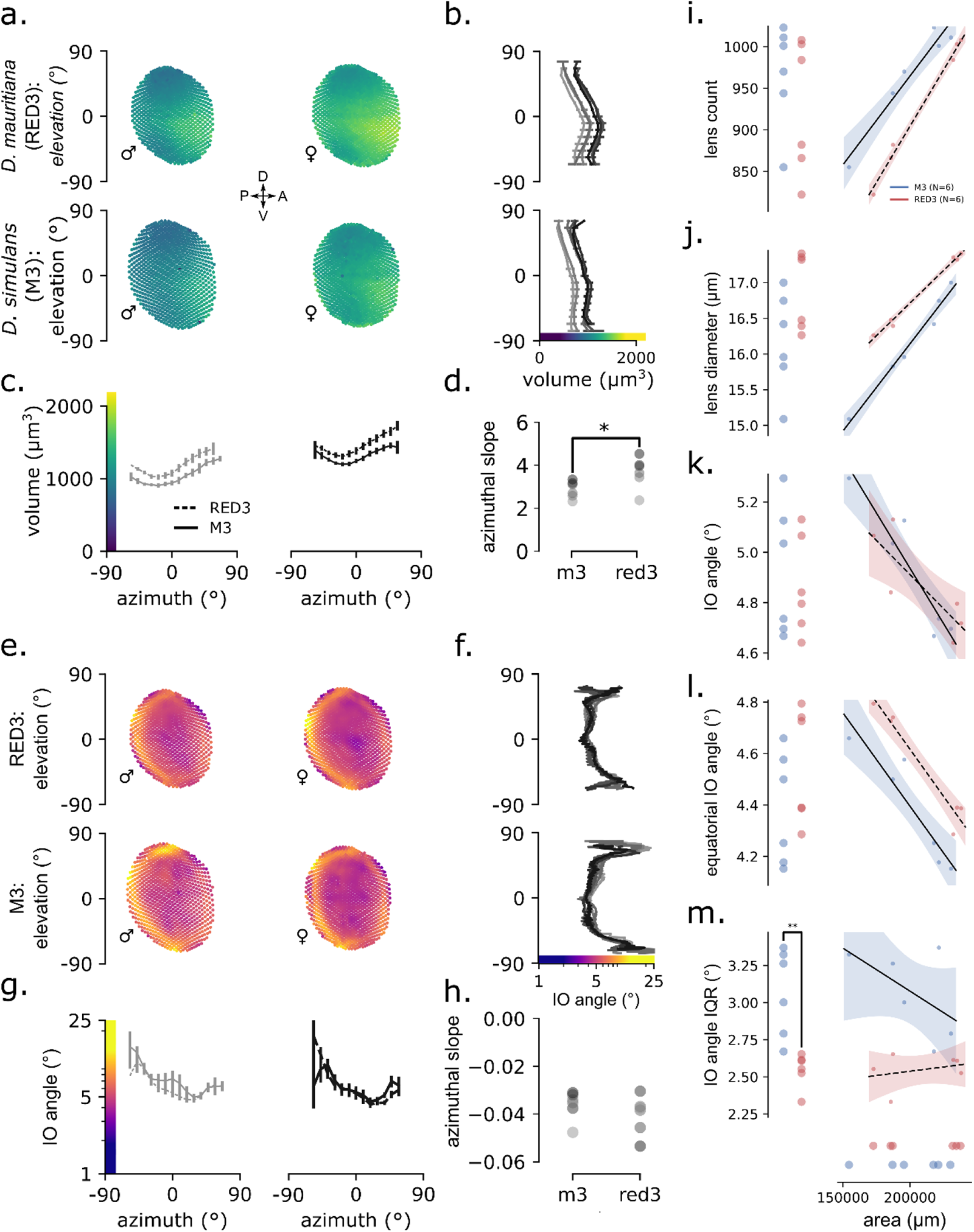
3D analysis of optic parameters in *D. simulans* and *D. mauritiana* eyes. **a–d.** Eye maps of ommatidial lens volumes from the smallest and largest eye from each of the two species: a male and female for each. Each dot of the scatterplot represents the location of an individual ommatidium in polar coordinates coloured by its 3D volume according to the colour scale indicated in the x- and y-axes of b and c. Line colours in b and c. and dot colours in d. indicate the fly’s rank in order of eye size per species, such that the darkest one is the largest eye of that species. The volume data is divided into 20 evenly spaced bins of elevation (**b.**) and azimuth (**c.**) with error bars indicating 3 times the standard error of the mean. **d.** Ordinary least squares were used to regress lens volume on azimuthal position to estimate and compare the azimuthal slope of lens volume between the two species. The resulting slope coefficients from those models are plotted. **e–h.** Eye maps of mean inter-ommatidial (IO) angle from the smallest and largest eye from each of the two species, plotted as in a-d. except for the elevation plot in f. The IO angle value used for each lens represents the average IO angle between that lens and all immediate neighbours. **f.** The same IO angle data from e. but sampling ommatidia from a narrow vertical band between 0±15° azimuth. Note that this is different from b., c., and g. because plotting the binned averages obfuscates the horizontal band of high acuity along the equator, likely due to the large range of IO angles along azimuth. **h.** Ordinary least squares was used to regress IO angle on azimuthal position as in d. All the eyes demonstrated negative azimuthal slopes, with no significant difference between species. **i–m.** Scatterplots of total lens count (**i.**), mean lens diameter (**j.**), median IO angle (**k.**), median equatorial IO angle (**l.**), and IO angle interquartile range (**m.**) plotted along the y-axes and their allometric relationship to the surface area of their eye along the x-axis. Lines in the 2D scatterplots represent the predicted mean and the bands represent the 95% CI of the mean based on ordinary least squares regression of each y variable on surface area. Note that simple group differences based on ANOVA are indicated in the left margins with the following key: * = p<.05, ** = p<.01, and ***=p<.001.

This area projects roughly onto the visual horizon during flight (Heisenberg and Wolf, 1988) and marks the region of the eye where rows of ommatidia initiated and grew during eye development, establishing a line of mirror symmetry about which rhabdomere arrangements flip vertically (Wolff and Ready, 1991). To compare the change in these parameters from the posterior to the anterior eye, we used ordinary least squares to fit an affine function of azimuth and compared the resulting slope parameters for each subject.

Both *D. simulans* M3 and *D. mauritiana* RED3 eyes have the largest lenses in the frontal visual field just below the eye equator (Fig. 3a). For both species, lens volume increases with elevation, peaks just below the eye equator, and then decreases steadily (Fig. 3b). In *D. mauritiana* RED3 this increase is more dramatically, starting at similar volumes at the dorsal and ventral extremes but increasing to larger maxima near the equator than *D. simulans* M3. Lens volume for both species decreases along elevation until a minimum around -45° and then increases, peaking at the anterior extreme (Fig. 3c). Moreover, in 5 of the 6 size-ordered pairs, *D. mauritiana* RED3 have significantly greater lens volumes than *D. simulans* M3 for every azimuthal bin (Suppl. Fig 5a). All eyes have positive azimuthal slopes, but the slope for *D. mauritiana* RED3 was significantly greater than *D. simulans* M3 (p=.043). This is consistent with measurements of lens diameter (Suppl. Fig 5b), cross-sectional area (Suppl. Fig 5c), and length (Suppl. Fig 6a), except that the azimuthal slope was significant for lens diameter (p=.047) but neither cross-sectional area (p=.18) nor length (p=.08). Overall, this means that *D. mauritiana* RED3 have larger, broader, longer, and wider-spread ommatidial lenses than *D. simulans* M3, which could improve sensitivity in general, and especially in the frontal visual field below the eye equator. This increase in ventral optical sensitivity is also more dramatic in *D. mauritiana* RED3 and is predicted to improve the detection of low-contrast objects below the visual horizon, such as rotting fruit or other oviposition sites.

### *D. mauritiana* RED3 and *D. simulans* M3 have higher spatial acuity along the eye equator

Inter-ommatidial (IO) angles are largest at the posterior and peripheral extremes, reaching a minimum around 45° azimuth along the eye equator (Fig. 3e and f). For both species, IO angle stays relatively constant—remaining between 4° and 6° from about -45° to 45° elevation— except for dramatic increases at the ventral and dorsal extremes and a region of smaller IO angles around the eye equator (Fig. 3f). *D. mauritiana* RED3 ranges less in IO angle than *D. simulans* M3, reaching smaller maxima in the top and bottom of the eye (≤15° versus ≤25°). For both species, IO angle decreases along azimuth from a maximum in the posterior extreme (≤15°) to a minimum around 45° azimuth (≥4°; Fig. 3g). We found no significant difference between species in the azimuthal profile or slope (Fig. 3g and h). Because spatial resolution is limited inversely by IO angle, maximum spatial resolution in both species is highest around 45° azimuth and 0° elevation, along the eye equator. This increase in equatorial spatial resolution might be an adaptation to terrain statistics of different habitats (Hughes, 1977; Currea et al., 2022), and due to the horizontal band of smaller ommatidial diameters at the eye equator formed during eye development (Ready, Hanson and Benzer, 1976; Kumar, 2012). Regardless, this would improve the resolution of small objects near visual horizon, a feature that would help in avoiding predators and locating oviposition sites.

### Eye allometry in *D. mauritiana* RED3 prioritizes contrast sensitivity more than *D. simulans* M3

In holometabolous insects, body size and the size of organs derived from imaginal discs depend on, and are proportional to, environmental factors like temperature and food availability during larval development (Shingleton et al., 2007; Shingleton, Mirth and Bates, 2008; Callier and Nijhout, 2013). In flies, larval feeding has been shown to affect eye size, ommatidia size, and ommatidia count (Currea, Smith and Theobald, 2018). As a result, variation in eye size and composition may reflect rearing differences. To address this, we modelled the scaling relationships between eye surface area and the following measurements: total lens count, mean lens diameter, median IO angle, median equatorial (elevation = 0±15°) IO angle, and IO angle interquartile range (Fig. 3i–m).

Eye surface area (*SA*) is an ideal reference for allometric scaling because it is proportional to the rate of light absorption of the entire eye and is approximately the number of ommatidial lenses (*N*) times the mean ommatidial lens area (*A*), *SA* ≈ *N* × *A*. Because the number of discernible brightness levels is proportional to lens area, *SA* is also proportional to the total number of images the eye can resolve, its spatial information capacity (Snyder, 1977). Using ordinary least squares, we regressed each measurement on the sum of eye area and a dummy-coded species variable (Suppl. Table 1). All models were a good fit, explaining a substantial proportion of the variance in SA plus the species variable (R^2^ = .65–.97, F = 8– 159, P ≤ .01).

Lens count and size had significant positive slope coefficients, such that larger flies have more and larger ommatidia in both species. However, pairwise tests for the *D. mauritiana* RED3 – *D. simulans* M3 difference had significant but opposite signs, indicating that *D. mauritiana* RED3 eyes have fewer but larger lenses and therefore lower ommatidial density than *D. simulans* M3. Despite these clear allometric differences, *D. simulans* M3 flies in this sample had generally smaller eyes but higher ommatidial density, resulting in similar lens counts between the two species. Conversely, the interquartile range (IQR) of lens diameters has a significant negative slope and a significant positive *D. mauritiana* RED3 – *D. simulans* M3 difference, implying that *D. mauritiana* RED3 lens diameters are more variable than *D. simulans* M3 after accounting for eye size. This is consistent with the lens volume eye maps discussed above, which found a greater range of lens sizes in *D. mauritiana* RED3 than *D. simulans* M3 along elevation, generally larger ommatidia for every azimuthal bin, and a greater azimuthal slope.

For both general and equatorial IO angles and comparison across both species, the slope coefficient was significant and negative, meaning that median angles scale inversely with eye size. However, the difference between species was only significant for median equatorial IO angles, such that *D. mauritiana* RED3 has significantly greater equatorial IO angles than *D. simulans* M3 after accounting for eye size. Because spatial resolution is inversely proportional to IO angle, *D. simulans* M3 has greater spatial resolution at the eye equator but similar resolution elsewhere. The IQR of IO angles had an insignificant slope coefficient and a significant but negative *D. mauritiana* RED3 – *D. simulans* M3 difference, meaning that *D. simulans* M3 have a greater range of IO angles. This is consistent with the IO angle eye maps above, which found a greater range in the elevation profiles of IO angle in *D. simulans* M3 (Fig. 3f). The increase in IO angles near the boundaries of the eye should effectively increase the FOV of the eye. Overall, these allometric relations suggest that *D. mauritiana* RED3 prioritize optical sensitivity more than *D. simulans* M3, which instead prioritize spatial resolution along the visual horizon and FOV at the peripheral extremes.

### *D. mauritiana* RED3 and *D. simulans* M3 optomotor responses trade off contrast sensitivity and spatiotemporal resolution

Our morphological analysis suggested that *D. simulans* M3 have higher spatial acuity due to smaller IO angles for equatorial ommatidia and *D. mauritiana* RED3 have higher optical sensitivity due to larger facet apertures, particularly in the central visual field just below the horizon. However, neural summation can recover sensitivity loss due to suboptimal optics by effectively sacrificing temporal or spatial resolution (Currea, Smith and Theobald, 2018). To measure the ethological implications of these optical differences, we used the flies’ optomotor response in a virtual reality flight simulator that allowed the presentation of different sinusoidal gratings moving to the left or right (Suppl. Fig. 7). Using a wingbeat analyser, we measured the flies’ steering effort in response to gratings of various contrasts, spatial frequencies, and temporal frequencies sorted randomly. Contrast sensitivity is defined as the reciprocal of the lowest contrast response, and both spatial and temporal acuity are defined by the maximum discernible frequency. Assuming that the interommatidial angle limits the maximal spatial sampling or resolution of the eye according to the Nyquist limit, such that the highest possible discernible frequency, *f_s_*, for a hexagonal lattice is given by the following equation: *f_s_* = 1/√3 * *Δɸ*^-1^. So, for every *f_s_*, there is a corresponding ideal IO angle, *Δɸ* = 1/√3 * *f_s_* ^-1^.

In the flight arena, *D. simulans* M3 and *D. mauritiana* RED3 traded off between higher contrast sensitivity and spatiotemporal tuning (Fig. 4). In accord with their larger ommatidia, *D. mauritiana* RED3 demonstrated higher contrast sensitivity (.14^-1^=7.4) than M3 (.27^-1^=3.7). Conversely, the spatial tuning curves demonstrate that *D. simulans* M3 has a higher spatial acuity (0.1 cpd) than *D. mauritiana* RED3 (0.08 cpd), implying smaller IO angles (∼5.8° versus ∼7.2°) consistent with smaller measured IO angles in the eye equator of *D. simulans* M3. *D. simulans* M3 also responded with greater strength around 0.04 cpd, likely supported by their wider peripheral IO angles and greater IQR. Lastly, *D. simulans* M3 demonstrated higher temporal acuity, 50 Hz, than *D. mauritiana* RED3, 20 Hz. Overall, this demonstrates sharper (higher spatial acuity) and faster vision (higher temporal acuity) in *D. simulans* M3 but a greater ability to compare brightness values (higher contrast sensitivity) in *D. mauritiana* RED3.

**Figure 4:**
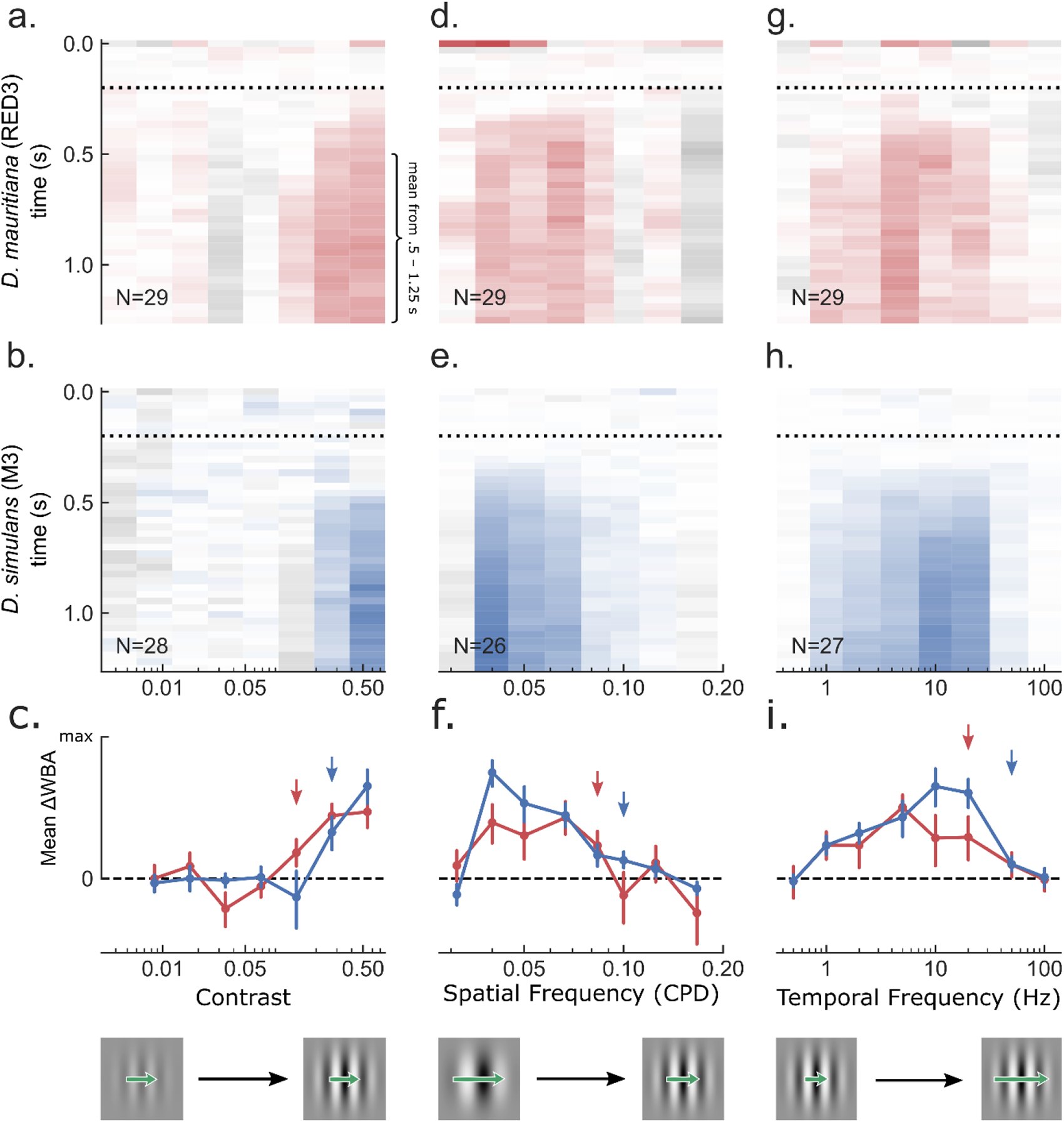
Behavioural measurement of *D. simulans* and *D. mauritiana* contrast sensitivity, spatial resolution, and temporal resolution. Gratings of various contrasts (**a**-**c**), spatial frequencies (**d**-**f**), and temporal frequencies (**g-i**) were presented to the flies in a rigid tether flight simulator equipped with a wingbeat analyzer. The gratings were filtered through a gaussian window and remained still for .2 s before moving to the left or right, indicated by the dotted line. For each subject, responses to leftward moving gratings were averaged with responses to the same grating moving rightward so that positive values represent mean steering in the direction of the grating (red or blue) and negative represents counter steering (gray). Mean normalized responses were baseline corrected, subtracting the mean response during the .1s before the onset of motion. Sample sizes are indicated in the bottom left corner of the colormaps. The images of gratings in the bottom of **c.**, **f.**, and **i.** are meant to give a sense of the change in the stimulus along the x-axis. Green arrows indicate the change in speed of the grating, ft/fs, which remains constant in the contrast experiment, decreases in the spatial frequency experiment, and increases in the temporal frequency one. **a**-**c.** As contrast increases, RED3 begins responding significantly at .14 (red arrow in **c.**) and M3 at .27 (blue arrow in **c.**). **d**-**f.** As spatial frequency increases and therefore rotational velocity decreases, mean responses decrease gradually until the Nyquist limit determined theoretically by the IO angle, reducing the contrast for higher frequencies as a result of aliasing. This limit differed between the two species, with RED3 responding significantly to spatial frequencies as high as .08 CPD (red arrow in **f.**) and M3 as high as .1 CPD (blue arrow in **f.**). **g**-**i.** As temporal frequency and therefore rotational velocity increases, mean responses increase until they reach the Nyquist limit determined by the temporal resolution of the optomotor response, reducing the contrast for higher frequencies. M3 demonstrated higher temporal acuity, responding significantly to frequencies as high as 50 Hz (blue arrow) while RED3 stopped at 20 Hz (blue arrow).

## Discussion

Variation in overall eye size, ommatidia number and facet area has been shown within and between *Drosophila* species by several research groups and differences in vision have been proposed based on these optical parameters (Posnien *et al*., 2012; Arif *et al*., 2013; Hilbrant *et al*., 2014; Keesey *et al*., 2019; Ramaekers *et al*., 2019; Gaspar *et al*., 2020). In this study the demonstrated allometries and regional specializations of *D. mauritiana* and *D. simulans* were found to differ in quantity but not quality: 1) maximum sensitivity in the central visual field below the horizon, with very similar elevation and azimuthal profiles; 2) maximum acuity along the visual horizon of the eye; and 3) improvements in optical sensitivity and spatial resolution for larger conspecifics. Future investigations of the developmental origins of these gradients and regional specializations in spatial resolution and optical sensitivity and how they may differ between flies will aid our understanding of how the astonishing diversity in insect eyes has evolved.

So far, our knowledge of how insects developmentally and evolutionarily balance the trade-off between ommatidia number (resolution) and ommatidia size (sensitivity) is very limited. The genetic basis of evolutionary differences in eye size has been difficult to determine, partly because ommatidia size and number seem to be genetically uncoupled and differences in these features polygenic. While differences in ommatidia size between *D. mauritiana* and *D. simulans* have been mapped to *orthodenticle* (Arif et al., 2013; Torres-Oliva et al., 2021), and a cis-regulatory region of *eyeless* has been shown to contribute to differences in eye size within *D. melanogaster* and between this species and *D. pseudoobscura* (Ramaekers et al., 2019), these changes do not explain the full extent of variation. Other genes involved in the regulation of cell proliferation and differentiation in developing eye imaginal discs are the most likely candidates to contribute to the diversification of eye size. For example, phosphoinositides including the *Drosophila* class I(A) PI 3-kinase Dp110 and its adaptor p60, the gap-junction protein *inx2* and the 40s ribosomal protein S6 kinases, have all been shown to alter ommatidia number and/or size (Montagne et al., 1999; Weinkove et al., 1999; Richard and Hoch, 2015; Janardan et al., 2020).

Very little comparative functional data is available to truly understand the impact of natural variation in eye structure on vision. Here we modelled and tested optical capacity in two *Drosophila* species – *D. mauritiana* and *D. simulans* – between two strains that had similar ommatidia number but significantly different ommatidia facet sizes, to assess whether predicted differences in contrast sensitivity, spatial resolution, and temporal resolution could be observed in behavioural experiments. In principle, larger facets could evolve to provide either better sensitivity (collecting more light in a similar amount of time) or better temporal resolution (collecting similar amounts of light over shorter times), or some combination. Indeed, we confirmed higher spatial and temporal acuity in *D. simulans* with smaller ommatidia, and improved contrast sensitivity in M3, with larger ommatidia. Whether and how these differences reflect meaningful adaptations to ecological differences remains to be explored, but the recapitulation of morphological divergence through behavioural paradigms is compelling.

We also identified substantial intraocular variation in lens volume, interommatidial spherical angles, facet shape, and lens diameter, as has been reported in other dipterans. While the optomotor experiments reported here tested global responses, future behavioural experiments might target different parts of the visual field to see whether this regional variation has a functional significance. Potentially increased spatial resolution at the equator of the eye, and in the anterior-ventral FOV, was predicted from our morphological data. Increased acuity at the horizon, combined with horizontally narrower facets at the centre of the eye, for example might enhance the detection of lateral optic flow. Likewise, the stronger anterior-frontal gradient in predicted resolution for females could have implications for the detection of oviposition sites.

While our analysis supports the use of 3D morphological data to predict optical capacity, many other factors are involved in information acquisition and processing in the insect eye that are not as easily accessible. Recent discovery of smooth and saccadic retinal muscle movement to improve perception of moving and stationary objects respectively (Hengstenberg, 1972; Fenk et al., 2022) as well as hyperacute vision via photomechanical photoreceptor contractions (microsaccades) (Hardie and Franze, 2012; Juusola et al., 2017; Kemppainen, Mansour, et al., 2022; Kemppainen, Scales, et al., 2022) have revealed much more sophisticated mechanisms are employed in *Drosophila* eyes to sample visual information. In particular, the spatial resolution of compound eyes can exceed the spatial Nyquist limit set by the IO angle due to brief, stereotyped photomechanical contractions (microsaccades) that sharpen and shift rhabdomere receptive fields, affording so called hyperacuity (Juusola *et al*., 2017). These contractions are optimal for processing brief bursts of light followed by periods of darkness to better match the refractory phase of rhabdomere microvilli (Juusola *et al*., 2017) and generally match the optical flow of forward translation (Kemppainen, Scales, *et al*., 2022). As a result, these phases of improved acuity apply to specific combinations of motion direction, duration, speed, and visual field region. Moreover, the advantages and magnitude of the photomechanical rhabdomere contractions are limited by IO angle (Kemppainen, Mansour, *et al*., 2022), so that the difference in IO angles measured here still infer an important difference in visual capacity.

The complexity of the visual system overall, incorporating mechanisms of neural summation and hyperacuity, further highlights the importance of using behavioural measurements of acuity and sensitivity and reinforces the conceptual distinction between optical and contrast sensitivity. Neural summation could have reversed these differences as it did for *D. mojavensis* due to darkness adaptation (Currea *et al*., 2022) or facultatively within *D. melanogaster* individuals in response to forward optical flow (Theobald, 2017). An assessment of the optics alone would have ignored the difference in temporal acuity and overestimated the difference in contrast sensitivity between *D. mauritiana* and *D. simulans* based on differences in optical sensitivity.

Aside from the functional aspect, the maintenance of eyes and the underlying complex neurocircuits are a metabolically expensive investment (Laughlin, De Ruyter Van Steveninck and Anderson, 1998). For example, comparison between photoreceptor information rates of larger and more active flies like the blowfly *Calliphora* with the smaller *D. melanogaster* showed a five times higher performance in *Calliphora* but at a ten times higher energetic cost (Niven, Anderson and Laughlin, 2007). The evolution of overall larger eyes with more and wider ommatidia and resulting increase in contrast sensitivity in *D. mauritiana* must therefore represent an economically viable investment aligned to their specific optical needs. The balance between sensory system requirements and energy efficiency has been observed in other fly species: The male housefly (*Musca domestica*) has a 60% higher bandwidth (measure of speed of response) in their contrast-coding R1-6 compared to females, allowing them to track these females in flight, whereas bandwidth decreases in male blowflies by 20% towards the back of the retina (Hornstein *et al*., 2000; Laughlin, 2001). It is therefore conceivable that absolute eye size is under stronger selection than ommatidia number or ommatidia size on their own, and at least to some extent independent of body size and other functional traits (Shearn and Garen, 1974; Bryant and Levinson, 1985; Cowley and Atchley, 1990). Evidence from *Drosophila* wings suggests that compensatory mechanisms guarantee a certain wing size if overall size deviates too much (McCabe, French and Partridge, 1997; Calboli, Gilchrist and Partridge, 2003). A similar mechanism could be at play in *D. simulans* where ommatidia number and size to be coordinated to maintain similar eye size across strains.

Insects play vital roles in various ecosystems, including pollination and decomposition. Climate change and the disappearance of ecological niches around the world highlights the need understand how they perceive and interact with their environment and vision is a primary sensory modality for many insects, shaping their behaviour, foraging strategies, and reproductive patterns. Our study demonstrates that even subtle differences in ommatidia size between closely related species can have a measurable effect on their vision. Therefore, comparative studies of natural variation in eye morphology and the consequences for vision across dipterans and beyond are needed to fully understand how the diversification of eye size, shape and function allowed insects to adapt to the vast range of ecological niches around the world.

## Methods

### Fly strains and husbandry

Multiple strains of *D. simulans* and *D. mauritiana* were used in this study (Gaspar et al., 2020; Suppl. Table 2). All stocks used were kept on standard yeast extract-sucrose medium at 25°C under a 12:12 hr dark/light cycle. For experiments, flies were reared at controlled, low density, achieved by transferring set numbers of males and females (typically between 10-20 of each sex) into fresh food containers to lay offspring. Adult offspring were removed soon after eclosion for experiments.

### Scanning Electron Microscopy

Fly heads were prepared and imaged as previously described (Gaspar et al., 2020). Briefly, heads were fixed in Bouin’s solution (Sigma-Aldrich) and dehydrated to 100% ethanol. For SEM imaging heads were critical point dried in a Tousimis 931.GL Critical Point Dryer and mounted onto sticky carbon tabs on SEM stubs, gold coated (10 nm) and imaged in a Hitachi S-3400N SEM with secondary electrons at 5kV.

### Morphological measurements

SEM images of eyes were analysed using FIJI/ImageJ (Schindelin *et al*., 2012). For each strain, 15 males and 15 females were measured. Ommatidia number was counted manually by using multi-point tool for one compound eye per individual (from side views of compound eyes). Ommatidia size and overall eye area were measured manually with the polygon selection tool. Frontal and central ommatidia area were measured for each eye with the polygon selection tool. The area of six central ommatidia was average to determine mean ommatidia (facet) size. Wing and tibia of the second leg of each fly were dissected in 70% ethanol and mounted in Hoyer’s solution, and cured overnight at 60°C. Wings and tibia were imaged at 5x (1.25x) magnification using Zeiss Axioplan microscope equipped with ProgRes MF cool camera (Jenaoptik). Wings and tibias size were measured using the line tool in Fiji/ImageJ.

### Synchrotron Radiation Tomography

Fly heads were prepared as described for SEM to 100% ethanol, then stained with 1% iodine and washed in ethanol. Fly heads were mounted in 20 µl pipette tips filled with 100% ethanol for synchrotron radiation X-ray tomography and scanned at the TOMCAT beamline of the Swiss Light Source (Paul Scherrer Institute, Switzerland (Stampanoni et al., 2006) and Diamond-Manchester Imaging Branchline I13-2 (Diamond Light Source, UK) (Rau et al., 2011; Peić et al., 2013) as previously described (Gaspar *et al*., 2020; Torres-Oliva *et al*., 2021).

### 3D Segmentation

The IMOD Software package (Kremer, Mastronarde and McIntosh, 1996) was used to generate cropped mrc stacks for 3D segmentation and analysis of the head tissue, lenses and optic lobes in Amira v.2019.2 (Thermo Fisher Scientific). Ommatidial lenses were segmented through threshold and separate objects tools. Lens sizes were analysed and colour-coded depending on size with the label analysis and sieve module.

### Morphometric analysis

Heads were tilted in the SEM to obtain flat images of frontal and central ommatidia for geometric morphometric analysis. The six corners of each facet were landmarked using the *digitize2D* function in the R package Geomorph (v.4.0.4) (Baken *et al*., 2021; Adams *et al*., 2023). Data were registered and Procrustes transformed using *procSym* function in the package Morpho (v.2.10) (Schlager, 2017), to account for reflection, before principal component analysis using the Geomorph package. Hierarchical clustering was performed and visualised using the factomineR (v.2.6) and factoextra (v.1.0.7) (Lê, Josse and Husson, 2008; Kassambara and Mundt, 2020) packages, using the option *nb.clusters = -1* to select the optimal number of clusters. The strain, sex, and positional identities of the resulting clusters were analysed by Chi squared in base R, and the contribution of the principal components to clustering was extracted from *desc.var* generated by the *HCPC* function for clustering. Plots were generated using ggplot2 (v.3.4.2) (Wickham, 2017).

### Statistical analysis

Plots and statistical analysis were carried out in RStudio Version 2023.03.0+386 using the Tidyverse suite of packages (Wickham *et al*., 2019). Where analysis required comparison between a length and an area, the length measurement was squared. Linear lines of fit were added to plots using geom_line(stat=“smooth”, method=lm). Correlation statistics were calculated using Pearsons correlation coefficient using stat_cor(method = “pearson”).

### ODA and Allometry

To approximate the optical performance of the two species, we processed CT stacks of six flies (three male and three female) from the RED3 strain of *D. mauritiana* and the M3 strain of *D. simulans.* This allowed us to apply the 3D ommatidia detecting algorithm (ODA-3D; Figure Suppl. Fig. 7a), a pipeline for automatically measuring a number of visual parameters for compound eyes (Currea *et al*., 2023). Each dataset was manually cleaned to generate binary images of only the corneal lenses. Then, the program fitted a cross-sectional surface through the coordinates of the lens cluster and projected these coordinates onto the cross-section, allowing a custom clustering algorithm to find ommatidia-like objects in the 2D projected images. Finally, the volume, diameter, cross-sectional area, length, and average IO angle of each lens was measured. Eye surface area was estimated as the sum of the lens areas based on the ODA-derived lens diameters. Allometric scaling relations were derived by regressing each of the measured visual parameters on eye surface area plus a constant. The resultant parameters of these models are found in Supplementary Table 1.

### Flight Arena

To measure the optomotor performance of *D. mauritiana* RED3 and *D. simulans* M3, we performed psychophysics in a rigid tether flight simulator equipped with a wingbeat analyzer (technical information can be found in (Currea, Smith and Theobald, 2018; Currea *et al*., 2022) and Suppl. Fig. 7b. Flies were glued to a thin tungsten rod and centred within an acrylic cube lined with rear-projection material (with 1/6 of the panels left open), immersing them in the projection surrounding 5/6 of their FOV (Suppl. Fig. 7c). An IR light casted the shadow of each wing onto photodiodes below the fly designed to output the amplitude of each wingbeat shadow as a 1000 Hz voltage signal. The difference between the left and right wingbeat amplitudes (ΔWBA) is proportional to yaw torque and indicates the fly’s steering effort. For instance, in Suppl. Figure 7d we plot the ΔWBA time series for an exemplary fly in response to 9 gratings of different contrast (corresponding to the line’s saturation) moving to the left or right (warm vs. cool hue). Note that the strength of the response is affected by contrast while the direction corresponds generally to the direction of motion. These responses were taken from (Currea *et al*., 2022), which used the same methods.

### Psychophysics

In the flight arena, flies viewed gratings of various contrasts (Fig. 4a-c, with examples at the bottom of c.), spatial frequencies (Fig. 4d-f), and temporal frequencies (Fig. 4g-i) to measure the functional consequences of their optical differences. The gratings were filtered through a gaussian window and remained still for .2 s before moving to the left or right, indicated by the dotted lines in Fig. 4. For each subject, responses to leftward moving gratings were 1) averaged with responses to the same grating moving rightward, 2) baseline corrected, subtracting the mean response during the .1 s before the onset of motion, and 3) normalized to the maximum mean response per fly so that positive values represent mean steering in the direction of the grating, with a maximum of 1 (fully saturated red or blue) and negative represents countersteering (gray). These baseline-corrected normalized responses were averaged across each group to make the colormaps in Figure 4. For each fly, an average of these normalized responses was taken from 0.5–1.25 s and used for plotting and comparing means in the bottom row of subplots in Figure 4.

Bootstrapping was used to test for a grating’s discernibility by estimating the standard error of the mean and 90% C.I.s for the mean response. We bootstrapped the means taken between .5 and 1.25 s 10,000 times at the subject level to generate empirical sampling distributions of the mean for each parameter value accounting for repeated measures. The 68% C.I. of each distribution was used as an approximation of the standard error (error bars in the bottom row of Figure 4) and the lower bound of the 90% C.I. was used to test for positive significance with a two-tailed alpha of .1 or one-tailed alpha of .05. Contrast sensitivity was defined as the reciprocal of the lowest discernible contrast and spatial and temporal acuity were defined by the highest discernible frequency.

## Supporting information

raw data eye size variation

## Supplement

**Supplementary Figure 1:**
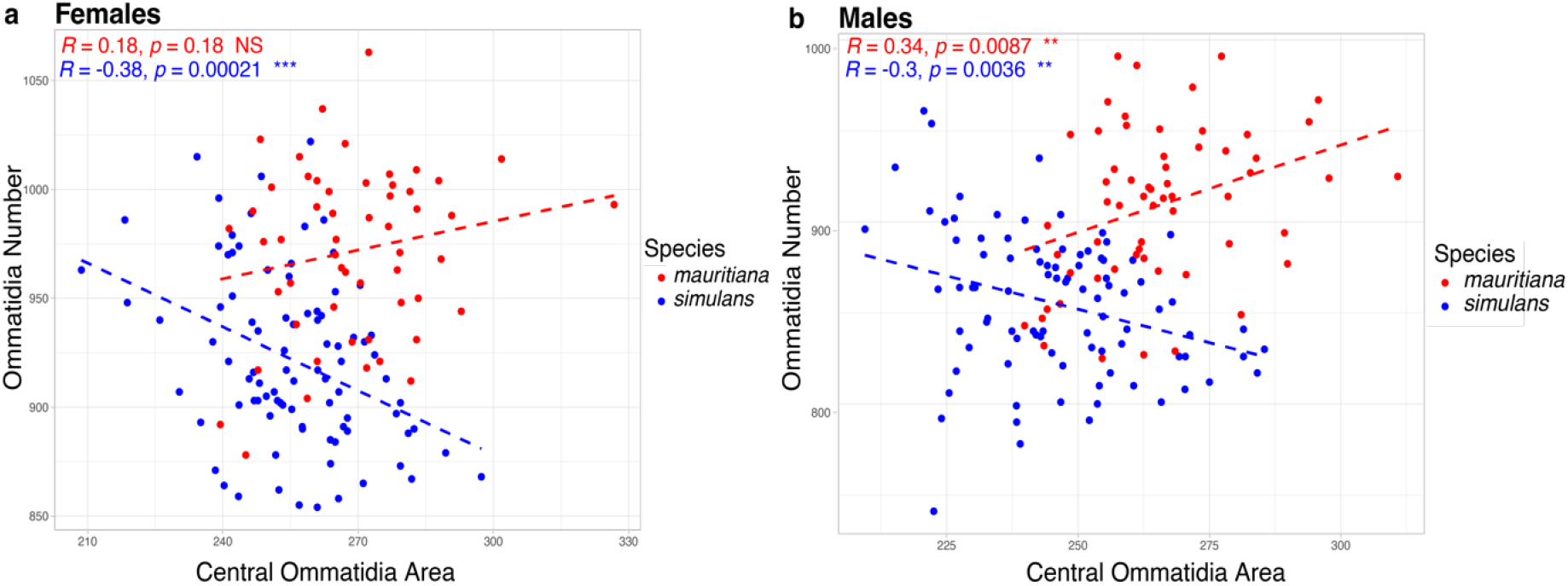
Correlation analysis for ommatidia number and ommatidia size *D. simulans* and *D. mauritiana* strains. Male and female *D. simulans* (blue) show a significant, negative correlation between ommatidia size and ommatidia number such that individuals with larger ommatidia tend to have less ommatidia overall. In *D. mauritiana* (red), males exhibit a significant positive correlation between ommatidia size and number, where the individuals with larger ommatidia also have a larger number of ommatidia.

**Supplementary Figure 2:**
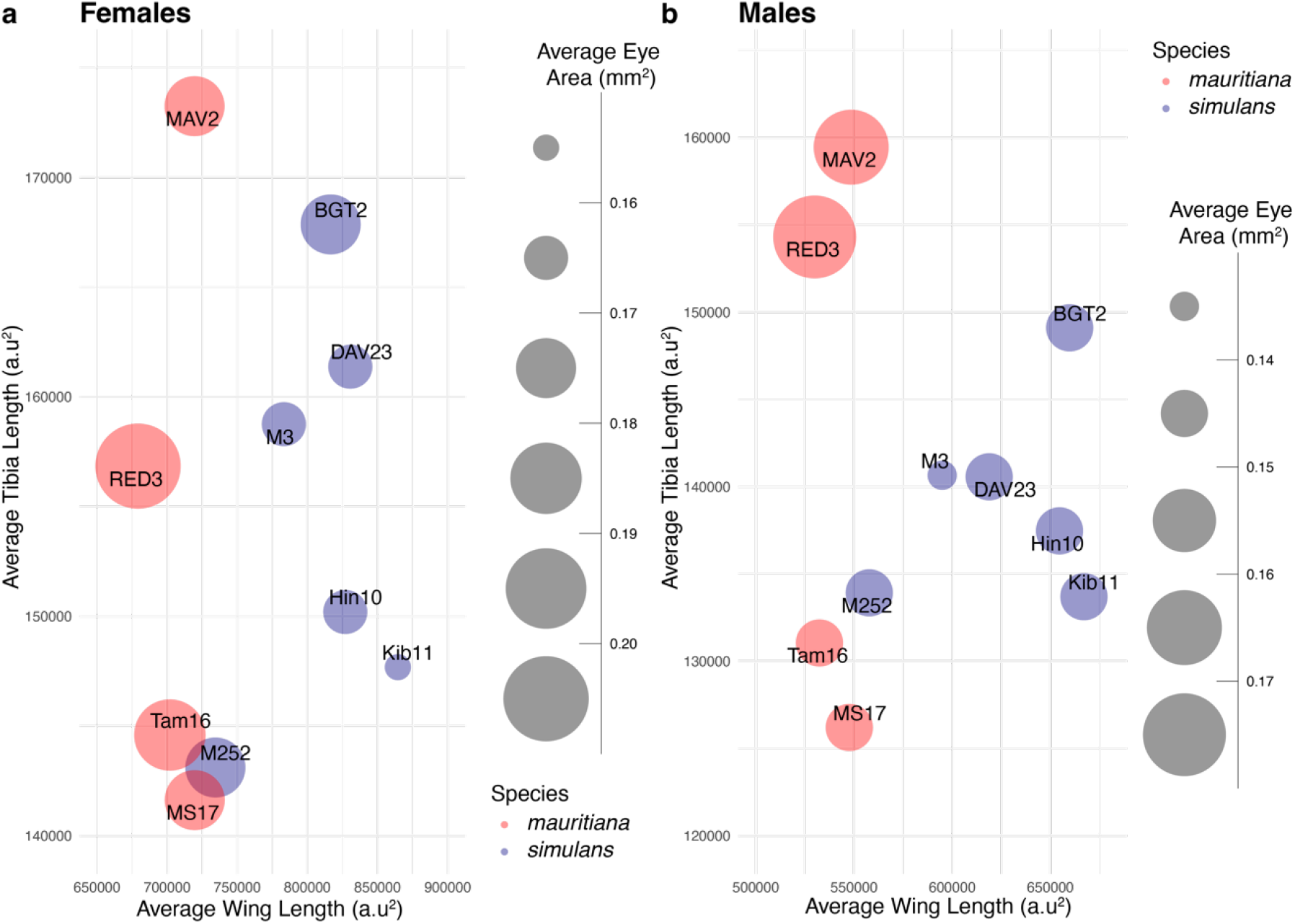
Variation in wing and tibia size across *D. mauritiana* (red) and *D. simulans* (blue) strains. Average eye size (circle area) of *D. simulans* (blue) and *D. mauritiana* (red) strains (circle labels) is plotted against wing vein and tibia lengths. *D. simulans* strains (blue) generally have larger wings but show some variation in tibia size whereas *D. mauritiana* strains generally have smaller wings and greater variation in tibia length.

**Supplementary Figure 3:**
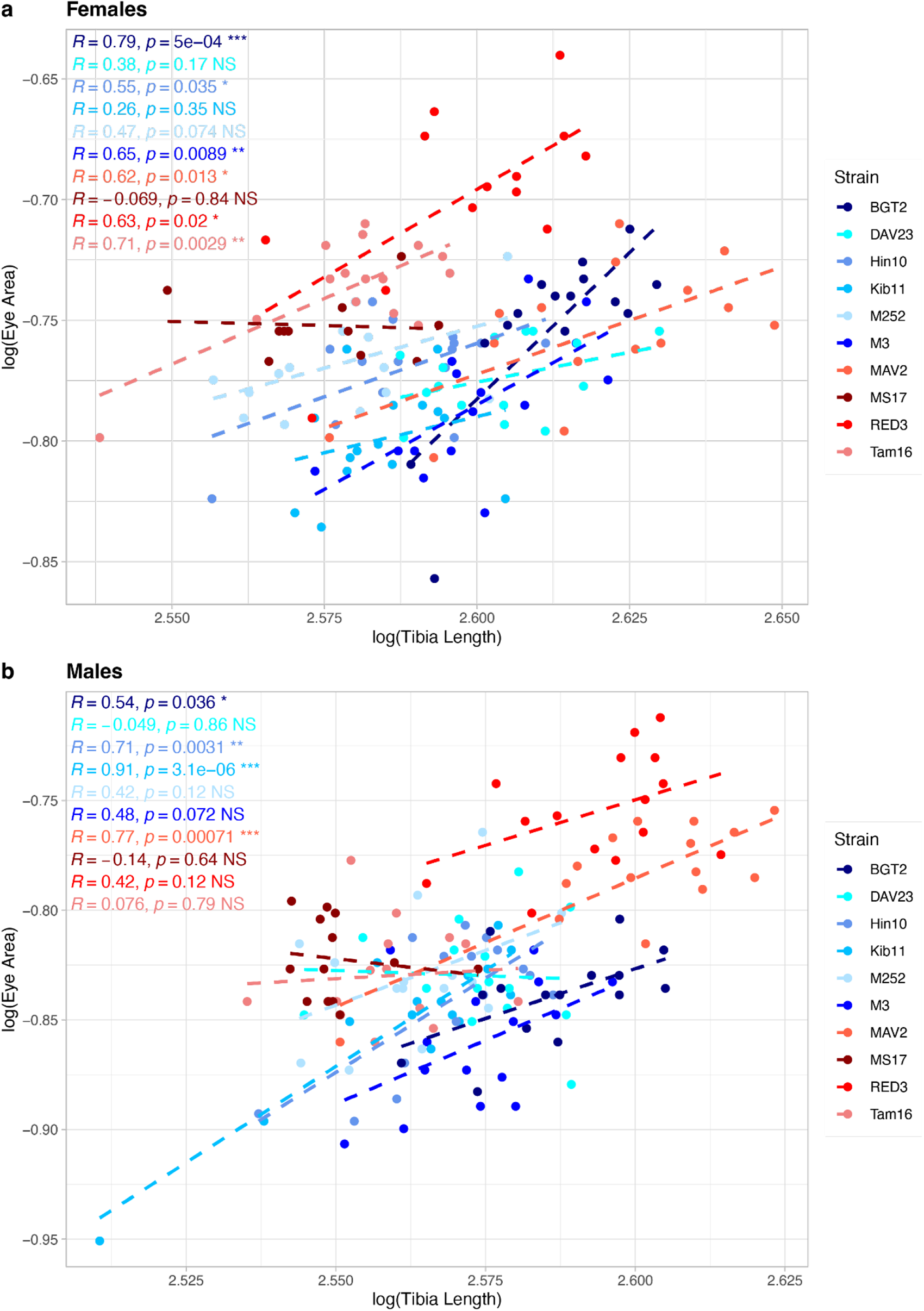
Correlation of 2^nd^ leg tibia size with eye size in *D. mauritiana* (red) and *D. simulans* (blue) strains. In males and females of both species only a subset of strains shows a significant positive correlation between tibia length and eye size.

**Supplementary Figure 4:**
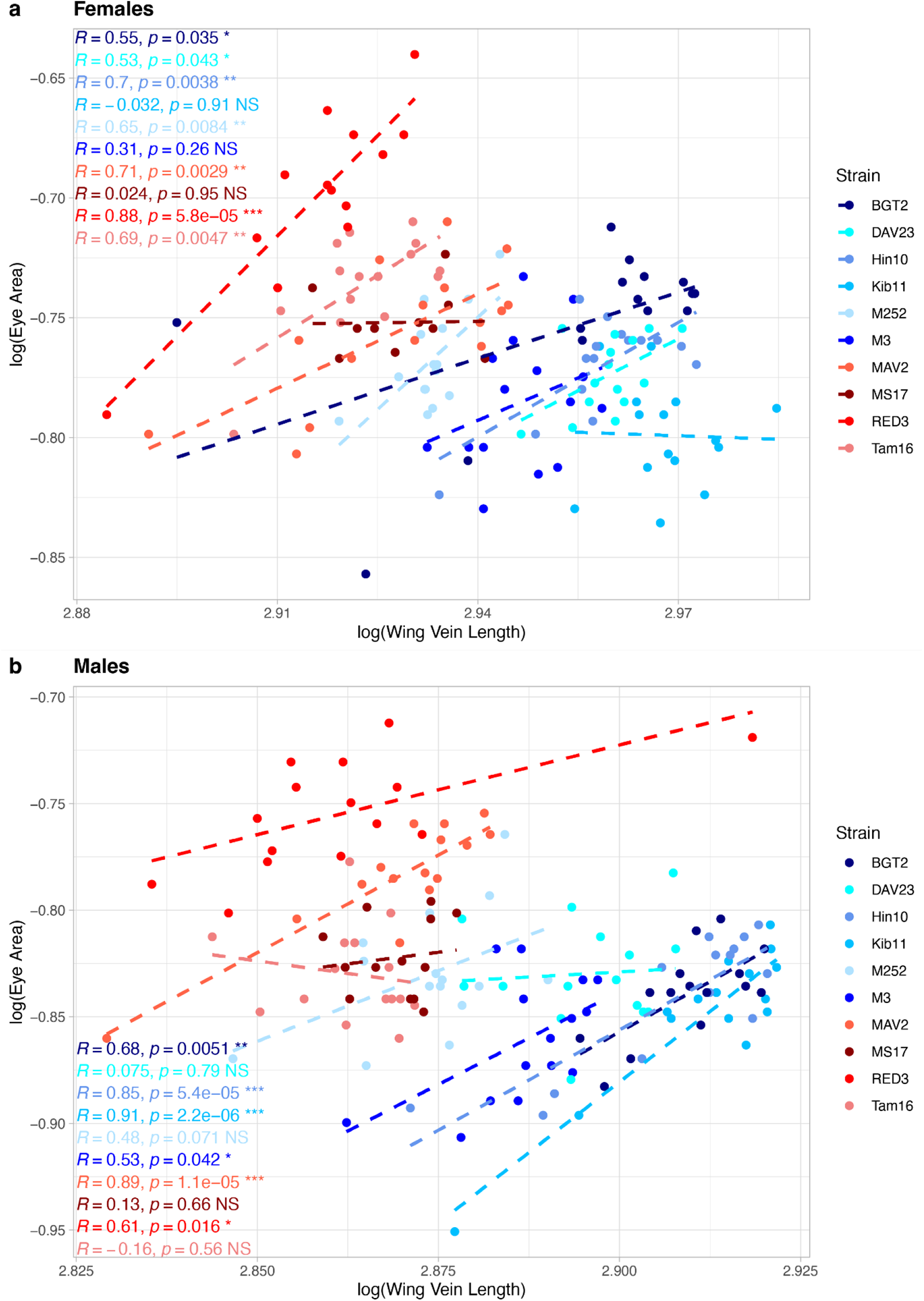
Correlation of wing size with eye size in *D. mauritiana* (red) and *D. simulans* (blue) strains. In males and females of both species only a subset of strains shows a significant positive correlation between wing vein length (a proxy for wing size) and eye size.

**Supplementary Figure 5:**
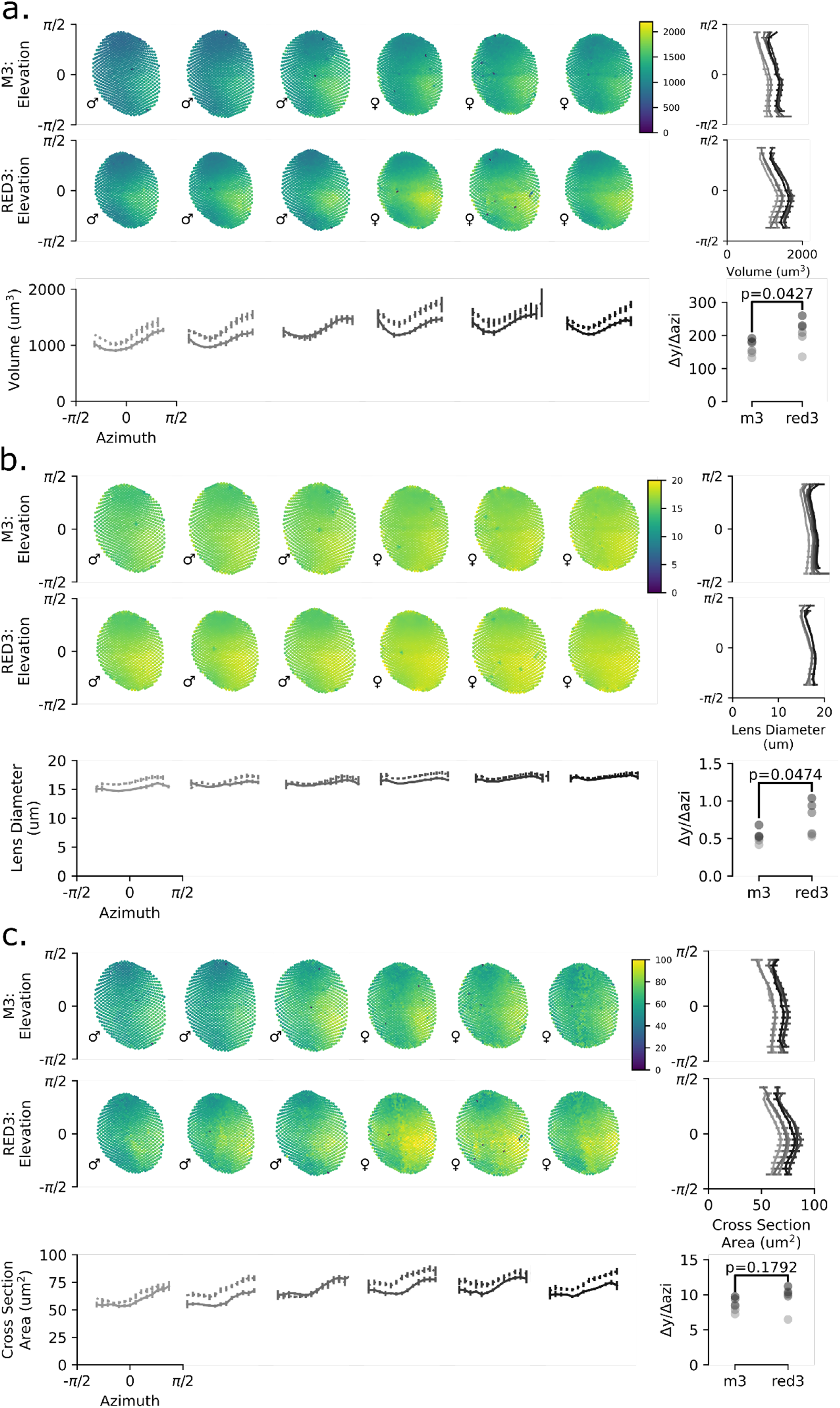
Eye maps of ommatidia measurements across eyes of *D. simulans* and *D. mauritiana*. **a–c.** Eye maps of ommatidial lens volumes (**a.**), lens diameters (**b.**), and cross-sectional areas (**c.**), with their elevation (right column of each panel) and azimuthal (bottom row of each) profiles, and their azimuthal slope (bottom right inset of each) from 6 flies from each of the two species, *D. mauritiana* (RED 3) and *D. simulans* (M3), 3 males and females for each. The eyes are sorted in order from smallest to largest eye surface area, which also resulted in ordering by sex because males are generally smaller. Each dot of the scatterplot represents the location of an individual ommatidium in polar coordinates coloured by its 3D volume according to the colour bars. Line colours in the azimuthal and elevation profiles and dot colours in the azimuthal slope plots indicate the fly’s rank in order of eye size per species, such that the darkest one is the largest eye of that species. Each outcome is divided into 20 evenly spaced bins of elevation (line plots to the right) and azimuth (line plots below) with error bars indicating 3 times the standard error of the mean of each bin. Ordinary least squares was used to regress each outcome on azimuthal position to estimate and compare the azimuthal slope between the two species. Scatterplots in the bottom right show the resulting slope coefficients from those models. Note that azimuth here is in radians but was converted to degrees for the plots Figure 3 and the calculation of the slope. **a.** Note that this presents the full dataset used in the elevation profiles of Figure 3b and the azimuthal slopes in Figure 3d.

**Supplementary Figure 6:**
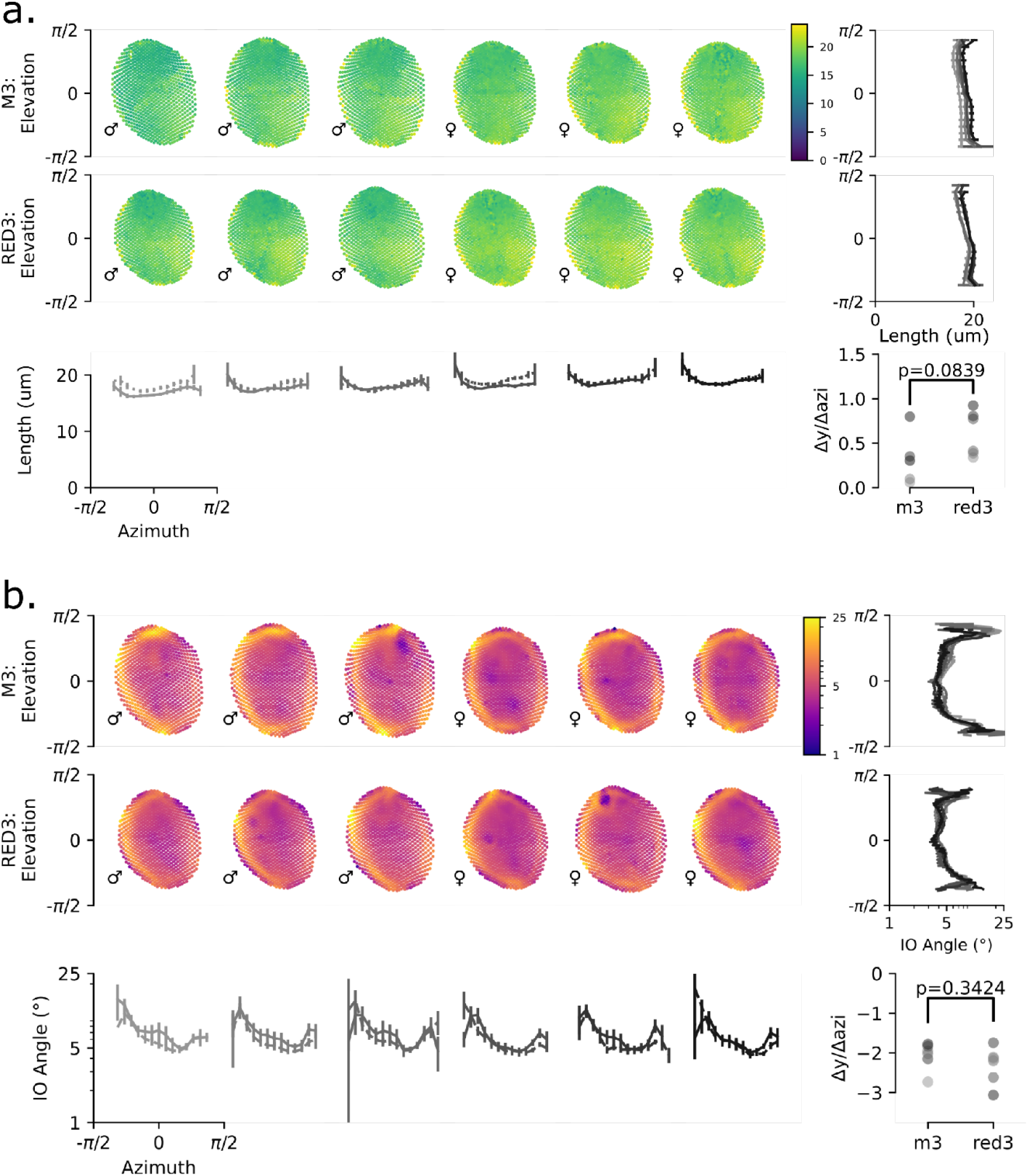
Eye maps of ommatidia lens length and IO across eyes of *D. simulans* and *D. mauritiana*. **a–b.** Eye maps of ommatidial lens length (**a.**) and IO angle (**b.**) as in Suppl. Figure 5, with their elevation (right column of each panel) and azimuthal (bottom row of each) profiles, and their azimuthal slope (bottom right inset of each) from 6 flies from each of the two species, *D. mauritiana* (RED 3) and *D. simulans* (M3), 3 males and females for each. The eyes are sorted in order from smallest to largest eye surface area, which also resulted in ordering by sex because males are generally smaller. Each dot of the scatterplot represents the location of an individual ommatidium in polar coordinates coloured by its 3D volume according to the colour bars. Line colours in the azimuthal and elevation profiles and dot colours in the azimuthal slope plots indicate the fly’s rank in order of eye size per species, such that the darkest one is the largest eye of that species. Each outcome is divided into 20 evenly spaced bins of elevation (line plots to the right) and azimuth (line plots below) with error bars indicating 3 times the standard error of the mean of each bin. Ordinary least squares was used to regress each outcome on azimuthal position to estimate and compare the azimuthal slope between the two species. Scatterplots in the bottom right show the resulting slope coefficients from those models. Note that azimuth here is in radians but was converted to degrees for the plots Figure 3 and the calculation of the slope. **b.** Note that, as in Figure 3, the elevation profile for IO angle was plotted differently because plotting the binned averages obfuscates the horizontal band of high acuity along the equator, likely due to the large range of IO angles along azimuth. This also presents the full dataset used in Figure 3f and the azimuthal slopes in Figure 3h.

**Supplementary Figure 7:**
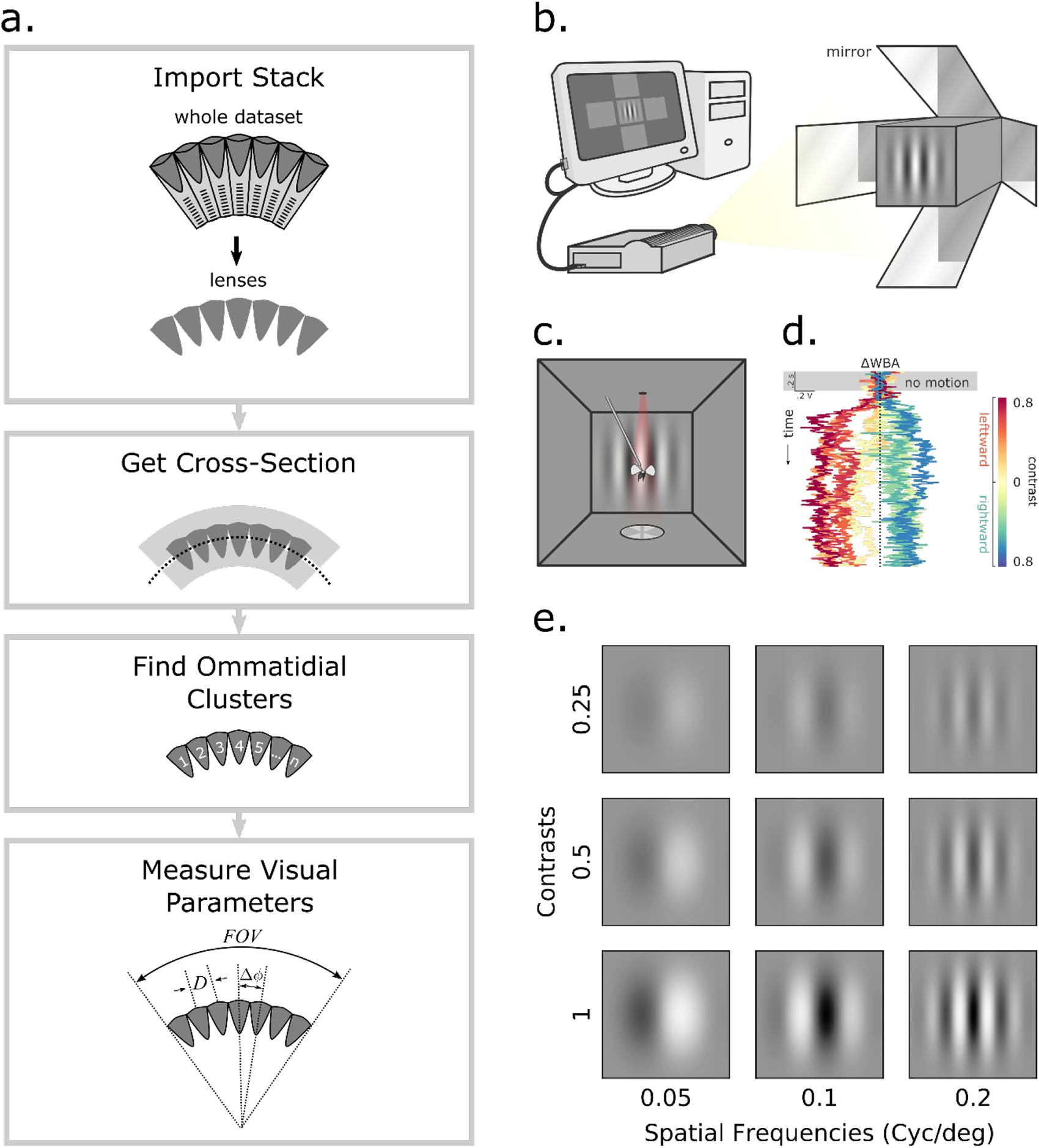
Workflow for modelling and testing vision in *Drosophila*. **a.** Optical performance was evaluated across the visual field of each eye by applying the ODA-3D, which requires first prefiltering the stack to just its corneal lenses to fit a cross-sectional surface, apply a clustering algorithm, and finally take optically relevant measurements for each lens. **b.** Optomotor performance was evaluated by a virtual reality flight simulator using an open-source computer graphics library, highspeed projector, and precisely positioned first-surface mirrors to project high resolution and contrast stimuli surrounding 5/6 of the flies’ FOV. **c.** Flies were glued to a thin tungsten rod and centred within an acrylic cube lined with rear-projection material immersing them in the projection as in b. An IR light casted the shadow of each wing onto photodiodes below the fly designed to output the amplitude of each wingbeat shadow as a 1000 Hz voltage signal. The difference between the left and right wingbeat amplitudes (ΔWBA) is proportional to yaw torque and indicates the fly’s steering effort. **d.** For instance, we plotted the ΔWBA time series for an exemplary fly in response to 9 gratings of different contrast (corresponding to the line’s saturation) moving to the left or right (warm vs. cool hue), drawn from (Currea *et al*., 2022). Notice that the strength of the response is partially dependent on contrast while the direction corresponds generally to the direction of motion. Leftward motion ΔWBA responses were averaged with the inverse of rightward motion ΔWBA responses to account for directional biases in our measurement. These averages were then normalized to the maximum mean response per fly and averaged across each group to make the colormaps in Figure 4. For each fly, an average of these normalized responses was taken from 0.5–1.25 s and used for plotting and comparing means in the bottom subplots (Figure 4 c, f, and i). These. **e.** Sinusoidal moving gratings were used because they are independently defined by a single orientation (leftward or rightward, for example), spatial frequency (x-axis), contrast (y-axis), and temporal frequency (the frequency of brightness change per pixel).

**Supplementary Table 1:** Parameters of the linear regression models of the allometries of optical parameters with respect to eye surface area. Each outcome was modelled as a linear combination of eye area and species (dummy-coded) with a constant intercept using ordinary least squares regression. The coefficient of determination (R^2^) and F-statistic are provided as measurements of goodness-of-fit with asterisks indicating the significance according to the key at the bottom of the table. The intercept and slope are the resulting coefficients of the regression model. *D. mauritiana* RED3 – *D. simulans* M3 is the pairwise difference of means after accounting for differences in eye size, such that values < 0 imply that *D. simulans* M3 values were greater than *D. mauritiana* RED3 relative to eye size. The significance of these statistics (F, intercept, slope, and *D. mauritiana* RED3 – *D. simulans* M3) is signified by the number of asterisks next to these values according to the key at the bottom of the table.

| outcome | $R^2$ | F | intercept | slope | RED3 – M3 |
| --- | --- | --- | --- | --- | --- |
| lens count | 0.96 | 110*** | 976*** | $2.4 \times 10^{-3}$*** | -58*** |
| lens diameters ($\mu\text{m}$) | 0.97 | 159*** | 16*** | $2.1 \times 10^{-5}$*** | 0.54*** |
| lens diameter IQR ($\mu\text{m}$) | 0.73 | 12** | 1.1*** | $-3.4 \times 10^{-6}$* | 0.33** |
| median IO angle ($^\circ$) | 0.78 | 16** | 4.9*** | $-7.1 \times 10^{-6}$*** | -0.009 |
| median equatorial IO angle ($^\circ$) | 0.92 | 53*** | 4.4*** | $-7.3 \times 10^{-6}$*** | 0.22*** |
| IO angle IQR ($^\circ$) | 0.65 | 8** | 3.1*** | $-2.2 \times 10^{-6}$ | -0.51** |
| **key:**<br>*: $p \leq .05$, **: $p \leq .01$, ***: $p \leq .001$ | | | | | |

**Supplementary Table 2:**
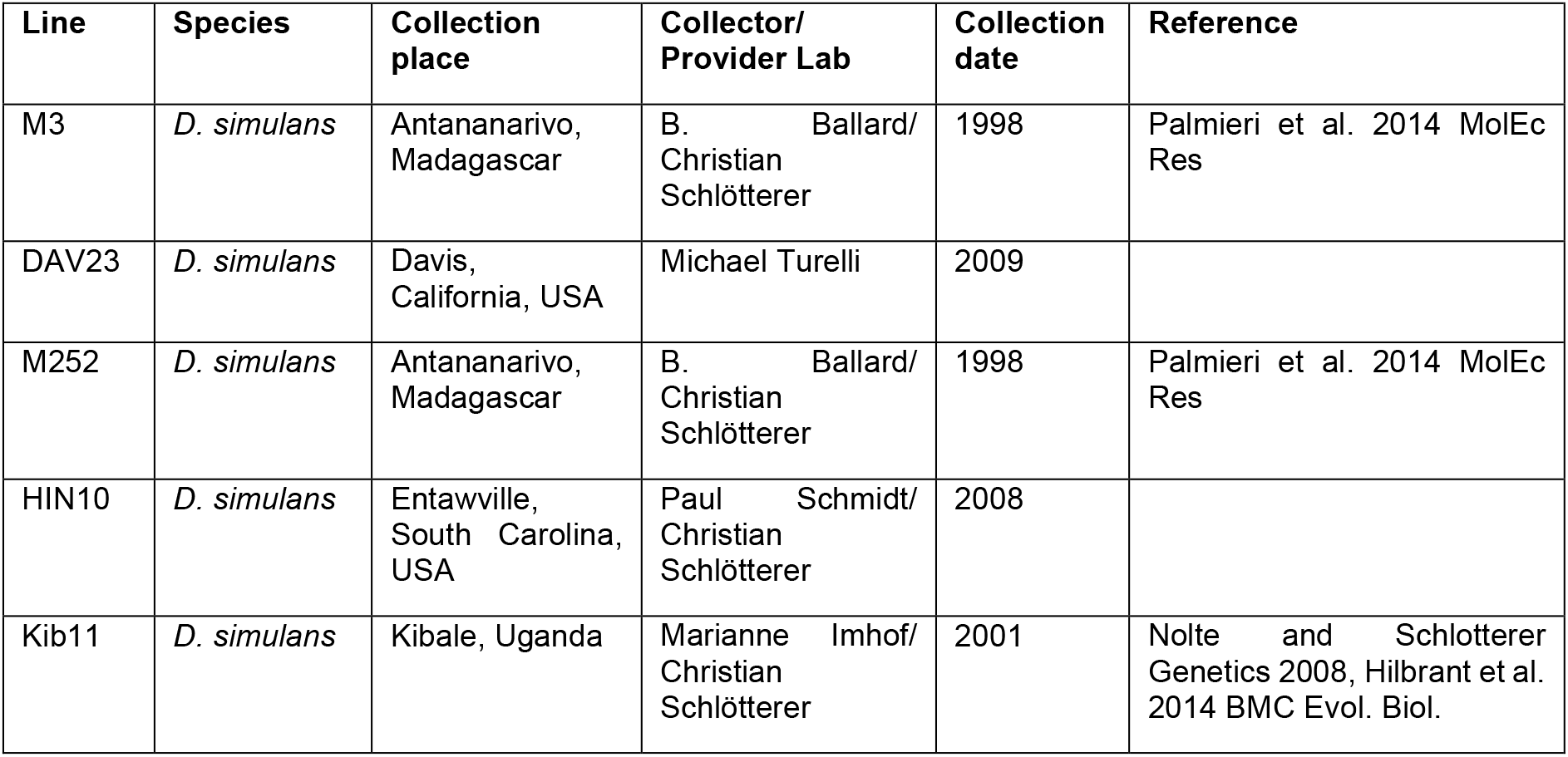
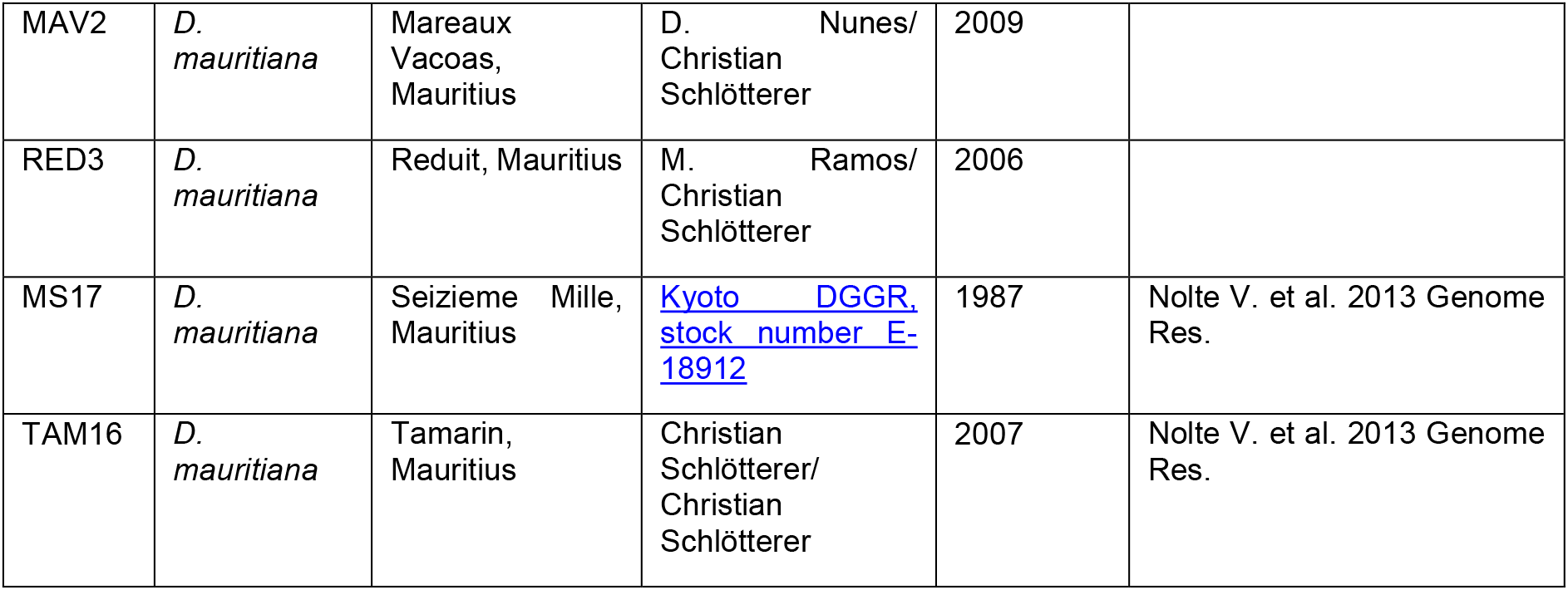
*Drosophila* strains used in this eye size survey.

| Line | Species | Collection place | Collector/ Provider Lab | Collection date | Reference |
| --- | --- | --- | --- | --- | --- |
| M3 | *D. simulans* | Antananarivo, Madagascar | B. Ballard/<br>Christian Schlötterer | 1998 | Palmieri et al. 2014 MolEc Res |
| DAV23 | *D. simulans* | Davis, California, USA | Michael Turelli | 2009 |  |
| M252 | *D. simulans* | Antananarivo, Madagascar | B. Ballard/<br>Christian Schlötterer | 1998 | Palmieri et al. 2014 MolEc Res |
| HIN10 | *D. simulans* | Entawville, South Carolina, USA | Paul Schmidt/<br>Christian Schlötterer | 2008 |  |
| Kib11 | *D. simulans* | Kibale, Uganda | Marianne Imhof/<br>Christian Schlötterer | 2001 | Nolte and Schlötterer Genetics 2008, Hilbrant et al. 2014 BMC Evol. Biol. |
| MAV2 | *D. mauritiana* | Mareaux Vacoas, Mauritius | D. Nunes/<br>Christian Schlötterer | 2009 |  |
| RED3 | *D. mauritiana* | Reduit, Mauritius | M. Ramos/<br>Christian Schlötterer | 2006 |  |
| MS17 | *D. mauritiana* | Seizieme Mille, Mauritius | Kyoto DGGR, stock number E-18912 | 1987 | Nolte V. et al. 2013 Genome Res. |
| TAM16 | *D. mauritiana* | Tamarin, Mauritius | Christian Schlötterer/<br>Christian Schlötterer | 2007 | Nolte V. et al. 2013 Genome Res. |

## Acknowledgements

This work was supported by a BBSRC (BB/T000317/1) grant to A.P.M. and M.K. and a grant from the National Science Foundation, IOS-1750833 to JT. We thank Christian Schlötterer for kindly providing *D. simulans* and *D. mauritiana* strains. SEM imaging was done in the Centre for Bioimaging at Oxford Brookes University. We acknowledge the Paul Scherrer Institut, Villigen, Switzerland for provision of synchrotron radiation beamtime at the TOMCAT beamline X02DA of the SLS. We acknowledge Diamond Light Source for time on Beamline I13-2 under Proposal MG25391.

